# Evolution in *Sinocyclocheilus* cavefish is marked by rate shifts, reversals and origin of novel traits

**DOI:** 10.1101/2020.04.12.038034

**Authors:** Ting-Ru Mao, Ye-Wei Liu, Madhava Meegaskumbura, Jian Yang, Gajaba Ellepola, Gayani Senevirathne, Cheng-Hai Fu, Joshua B. Gross, Marcio R. Pie

**Author notes:** equal contribution (Ting-Ru Mao, Ye-Wei Liu, Madhava Meegaskumbura).

## Abstract

Epitomized by the well-studied *Astyanax mexicanus*, cavefishes provide important model organisms to understand adaptations in response to divergent natural selection. However, the spectacular *Sinocyclocheilus* diversification of China, the most diverse cavefish clade in the world harboring nearly 75 species, demonstrate evolutionary convergence for many traits, yet remain poorly understood in terms of their morphological evolution. Here, using a broad sample of 49 species representative of this diversification, we analyze patterns of *Sinocylocheilus* evolution in a phylogenetic context. We categorized species into morphs based on eye-related condition: Blind, Micro-eyed (small-eyed), and Normal-eyed and we also considered three habitat types (Troglodytic – cave-restricted; Troglophilic – cave-associated; Surface – outside of caves). Geometric morphometric analyses show Normal-eyed morphs with fusiform shapes being segregated from Blind/Micro-eyed (Eye-regressed) morphs with deeper bodies along the first principal component (“PC”) axis. The second PC axis accounts for shape complexity related to the presence of horns. Ancestral character reconstructions of morphs suggest at least three independent origins of Blind morphs, each with different levels of modification in relation to the typical morphology of ancestral Normal-eyed morphs. Interestingly, only some Blind or Micro-eyed morphs bear horns and they are restricted to a single clade (Clade B) and arising from a Troglodytic ancestral species. Our geophylogeny shows an east-to-west diversification spanning the Pliocene and the Pleistocene, with Troglodytic species dominating karstic subterranean habitats of the plains whereas predominantly Surface species inhabit streams and pools in hills to the west (perhaps due to the scarcity of caves). Integration of morphology, phylogeny and geography suggests *Sinocyclocheilus* are pre-adapted for cave dwelling. Analyses of evolutionary rates suggest that lineages leading to Blind morphs were characterized by significant rate shifts, such as a slowdown in body size evolution and a 3.3 to 12.5 fold increase in the evolutionary rate of eye regression. Furthermore, body size and eye size have undergone reversals, but horns have not, a trait that seem to require substantial evolutionary time to form. These results, compared to the *Astyanax* model system, indicate *Sinocyclocheilus* fishes demonstrate extraordinary morphological diversity and variation, offering an invaluable model system to explore evolutionary novelty.

## INTRODUCTION

Due to the absence of light, stable mean temperatures, absence of primary productivity, and paucity of dissolved oxygen, subterranean habitats are among the most challenging environments for life on earth (Ginet and Decou, 1977, Camacho, 1992). From surface-dwelling ancestral species, cavefish have secondarily adapted to live in cave systems, often demonstrating a remarkable array of morphological and behavioral adaptations (Soares and Niemiller, 2013; Yoshizawa, 2015). These involve enhanced sensation, and also dispensing of traits that incur a developmental or energetic cost. Cavefish species can be divided into two forms: troglophiles are closely associated with caves, but do not entirely depend on them, and troglobites are obligate cave dwellers (Dowling et al., 2002; Jingcheng and Weicheng, 2015). Troglobite fish *may* harbor special adaptations, such as complete eye loss, loss of pigmentation, changes in cranial symmetry, proliferation of neuromast sensory organs, development of horns, and in some species flat, hollow heads (Strecker et al., 2004; Zhao, 2006; Gross et al., 2008). Despite the ∼200 cavefish species from across the world, large diversification of cavefishes is rare. However, one extensive diversification occurs in *Sinocyclocheilus*, a monophyletic group of cyprinid fishes endemic to China, which allows a robust analysis of trait evolution relative to troglomorphism in a phylogenetic context.

The specialized traits cavefish bear have led them to be developed as models of evolution especially with respect to adaptations to novel environments and evolutionary convergence (Culver et al., 1995; Dowling et al., 2002; Jeffery, 2001; Li et al., 2008; Strecker et al., 2004; Yang et al., 2016). A lion’s share of knowledge on evolution and development in cavefishes has come from *Astyanax mexicanus* (Mexican tetra), a species with both surface-dwelling (pigmented and eyed) and cave-dwelling morphs (depigmented and blind), which can readily interbreed (Borowsky, 2008). In contrast to this well-studied model system, *Sinocyclocheilus* species not only include blind and normal-eyed morphs (Lan et al., 2013), but demonstrate a continuum from blind to normal-eyed species. Indeed, members of the *Sinocyclocheilus* genus display remarkable morphological evolution with divergent cave-dwelling, cave-associated, and surface-dwelling species.

*Sinocyclocheilus* species are thought to have shared a common ancestor in the late Miocene, undergoing a spectacular diversification spanning the Pliocene and Pleistocene across the southwestern parts of China’s 620,000 km^2^ of karst habitats (Huang et al., 2008), with nearly 75 extant species (Jiang et al., 2019). This resulted in an adaptive diversification into subterranean refugia traversing the intersection of the Guizhou, Guangxi and Yunnan provinces around the time of the uplifting of Tibetan/Guizhou plateau (Li et al., 2008).

One of the most striking forms of cave adaptation in *Sinocyclocheilus* is variation in eye morphology, categorized often into three morphs (Zhao and Zhang, 2009), ranging from Normal-eyed to Micro-eyed (small-eyed) to Blind species. Of all Chinese hypogean fishes, 56 species show troglomorphism such as reduction and/or loss of eyes, pigmentation, and the gas bladder. Presence of a horn-like structure and hyper-development of the dorsal protuberance (similar to a humpback whale) are two additional unique characters to certain Chinese hypogean species (Romero et al., 2009). These dramatic adaptations to cave life are reflected in the unique morphology of these fish.

The morphology of *Sinocyclocheilus* is most likely attributed to their habitat and local adaptations, however the precise function of certain morphologies (e.g., their horns and humps) remains unknown (Ma and Zhao, 2012). For instance, many blind species are obligate cave dwellers that have the ability to navigate along cave walls, cave-bottoms and within narrow passages (Yoshizawa, 2015). Yet, others are open water species that navigate in the manner of typical fish. There are also intermediate forms between these two principal morphs (Zhao and Zhang, 2009). However, the morphology of these fishes is so extreme that substantial variation in morphology is evident even within blind, intermediate and the open water species.

The pattern of body shape evolution in *Sinocyclocheilus* is a conundrum, which has not been addressed in an evolutionary context. Here, we explore key patterns of morphological evolution in these fishes, and demonstrate that the evolution in *Sinocyclocheilus* has been associated with significant rate changes and trait reversals across their phylogenetic history.

## MATERIALS & METHODS

### Phylogeny estimation

We compiled sequence data from GenBank for two mtDNA fragments (*NADH4* and *cytb*) of 39 *Sinocyclocheilus* species, and the outgroup species *Linichthys laticeps* (Cyprinidae). In addition, we generated sequence data for the *cytb* gene fragment of ten additional *Sinocyclocheilus* species (Table S1). For these species, total genomic DNA was extracted using the DNeasy Blood and Tissue Kit (Qiagen Inc., Valencia, CA) following the manufacturer’s protocols. DNA was amplified in 25-µL volume reactions: 3 mM MgCl_2_, 0.4 mM of dNTP, 1X buffer, 0.06 U of Taq DNA Polymerase, 2 mM of each primer. Thermocycling conditions included an initial step at 94 °C for 3 min, followed by 35 cycles at 45 s at 94 °C, 1 min at 46-50 °C and 45 s at 48-56 °C, and a final step at 72 °C for 5 min. PCR products were electrophoresed in a 1.5% agarose gel, stained with ethidium bromide and visualized under UV light. Successfully amplified products were purified using MicroconTM Centrifugal Filter Units (Millipore, Billerica, MA, U.S.A.). Sequencing reactions were carried out in 10 µl solutions including the following final concentrations: 5 ng/µl of template DNA, 0.5 µl of Big DyeTM (Applied Biosystems Inc., Foster City, CA, U.S.A.), 0.2 µM of each primer and 0.1X of reaction buffer. The final product was purified using SephadexTM G-50 (GE Healthcare Bio-Sciences AB, Uppsala, Sweden) for sequencing. Forward and reverse strands were reconciled using Staden v.1.6.0 (Staden, 1996). Sequences from both genes were concatenated and aligned unambiguously using ClustalW (Thompson et al., 2003), as implemented in MEGA v. 6.0 (Tamura et al., 2013), for a total alignment length of 2155 bp. We used JModelTest v.2.1.10 (Santorum et al., 2014) to determine the best models of evolution for each fragment, which were implemented in BEAST v.1.10.4 (Drummond and Rambaut, 2007) as a partitioned analysis to estimate the phylogenetic relationships and relative divergence times within *Sinocyclocheilus*. We calibrated the tree using the relative time period which diversification of *Sinocyclocheilus* initiated around 11.31 Mya, a reference point obtained from (Chen et al., 2018). We used a strict molecular clock and a calibrated Yule tree prior, as well as a GTR+I+G for each partition and ran the analysis for 20 million generations using the Cipres Science Gateway Server (Miller et al., 2010). Convergence was assessed by inspecting the log-output file in Tracer v.1.6 (Drummond et al., 2012), and by ensuring ESS values were above 200. The first 10% of the trees were discarded, and the post burn-in trees were used to infer the maximum clade credibility tree using TreeAnnotator v.1.4.4 (Rambaut and Drummond, 2019). The maximum clade credibility tree, as well as a set of 1000 post burn-in topologies, were retained for further analyses (see below).

### Morphometric data acquisition and analyses

We assembled a database of images of referenced specimens of *Sinocyclocheilus* from scaled photographs, which was complemented by species that we photographed (Table S2). The final dataset included 90 images (54 from species descriptions and catalogues and 36 from photographs by the authors) for 50 species, which included 70% of the total number of described species for the genus. These images were used for geometric morphometrics analyses, which were based on 15 landmarks (See Fig. 1A) and 180 sliding semi-landmarks, obtained using tpsDig v. 2.16 (Rohlf, 2010). Semi-landmarks were collected as curves outlining the body. These data were subsequently reduced to equidistant landmarks, and defined as semi-landmarks using tpsUtil v. 1.46 (Rohlf, 2010). We then slid the landmarks using the bending energy method (Gunz & Mitteroecker, 2013) implemented in geomorph v.3.2.0 (Adams & Otárola-Castillo, 2013). The landmark coordinates were aligned using a generalized Procrustes superimposition analysis (Adams et al., 2013), and a principal component analysis (PCA) was used to evaluate shape variation within the sample.

**Fig. 1.**
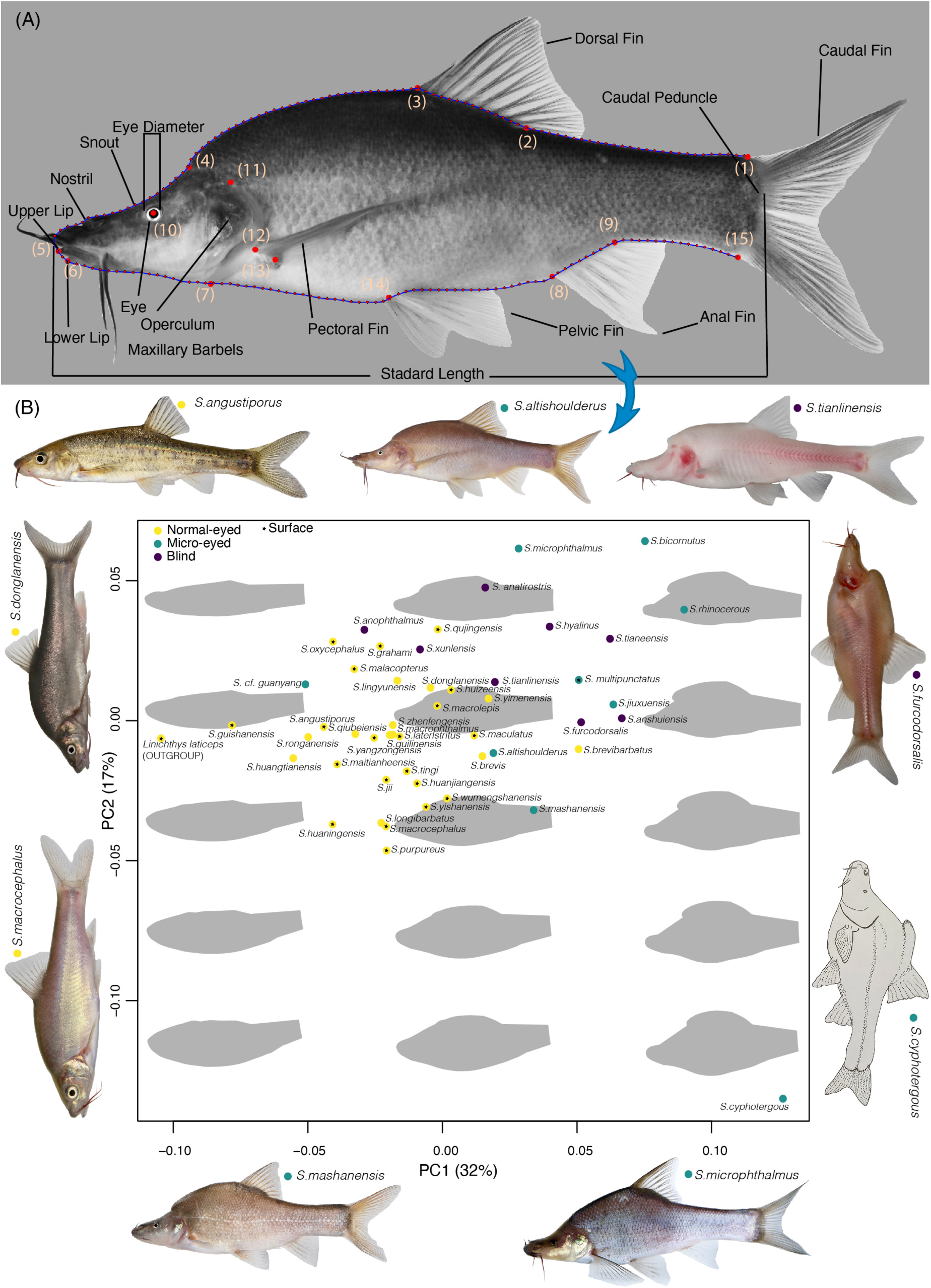
**(A) A specimen of *Sinocyclocheilus altishoulderus* indicating the position of 15 landmarks (red: larger points indicated by numbers 1-15) and 180 semi-landmarks (red: smaller point) used for the calculation of Procrustes coordinates and traditional linear measurements (SL: standard Length, ED: eye diameter and sED: standardized eye diameter) used in the geometric morphometric analyses**. (B) **PCA showing the variation in body shape of the genus *Sinocyclocheilus* traced with eye morphology and habitat occupation**. PC1 and PC2 accounts for 32% and 17% of the variance respectively. A shift from the fusiform shape of the Normal-eyed surface forms to a more “boxy” form of the Micro-eyed and Blind forms is evident.

Multiple images for the same species were used to obtain the landmarks and the mean of their Procrustes coordinates were calculated to be used in later analyses. We also obtained traditional linear measurements, namely standard length (SL), eye diameter (ED) and standardized eye diameter (sED, calculated as the ratio between ED and SL).

### Morphological and habitat evolution

Since shape variation in *Sinocyclocheilus* cavefishes occurs mostly in the anterior end of the fish and as one of the major features leading to this is eye-related, we considered the absence or the size of the eye (when present) as a proxy for the categorization of morphs. Since the eye size has an allometric association with body size, we used the standardized eye diameter (sED) in placing them into three morphological categories: Blind (eye absent); Micro-eyed (0.0 – 3.0mm); Normal-eyed (< 3.0mm) respectively. For ease of discussion, we considered Blind and Micro-eyed together as Regressed-eyed; Micro-eyed and Normal-eyed together as Eyed species.

*Sinocyclocheilus* were also categorized based on their habitat as Troglodytic, Troglophilic, and Surface species. Troglodytic species live in an obligatory association with caves, and are not sampled outside of caves. Caves, as meant here represent roofed-caves, submerged caves, and subterranean waterways that form windows intermittently with the surface. Troglophilic species live in a close association with caves and are sampled both in the vicinity of cave entrances and within caves. Finally, Surface species are found in habitats even when a cave is not found in close proximity and live in normal streams ponds and lakes as typical fish do, but they could venture into caves (underground water bodies) during unfavorable periods, when water is only available in caves. It should be noted here that this categorization is strictly habitat based and not morphology based (for instance, there are Normal-eyed species that are Troglodytic, Troglophilic or Surface). These habitat associations are based on published literature (Zhao and Zhang, 2009; Romero et al., 2009) and personal observations as outlined in Table S1.

To infer the number and timing of evolutionary shifts within eye-related morphs, horn distribution and the habitat type, we used stochastic character mapping (Nielsen, 2002; Huelsenbeck et al. 2003), as implemented in the make.simmap function in phytools. On each of the 1000 post burn-in trees obtained from BEAST, we used stochastic character mapping to generate 100 potential histories. This approach therefore considers uncertainty both in the evolutionary history of the traits as well as in the inferred topology of the phylogeny.

The landmark coordinates obtained were aligned using a generalized Procrustes superimposition analysis (Adams et al., 2013), and a principal component analysis (PCA) was used to explore shape variation within the sample. In addition, we described the eye-related morphological variation in *Sinocyclocheilus* by estimating ancestral states of SL and ED and visualizing them using traitgrams (Evans et al., 2009) as implemented in the phenogram function in phytools 0.7.20 (Revell, 2012) using the maximum clade credibility tree. We also visualized the evolution of both traits simultaneously using a phylomorphospace (a projection of the tree into morphospace, *sensu* Sidlauskas, 2008) using the phylomorphospace function in phytools.

### Evolutionary rate variation in eye related morphs

We tested whether the evolutionary rates of the studied continuous traits (SL, ED, sED) are significantly different in different morphs. We used 100 potential trait histories from stochastic character mapping and then fit two alternative models of evolution on each studied trait, one that fixes the rate of evolution to be identical between morphs against an alternative model in which the morphs have separate rates. We calculated the Akaike Information Criterion for small sample size (AICc) from the maximum likelihood estimate on each tree using the brownie.lite function in phytools. Finally, if a multi-rate model provided a better fit to the data, we calculated model-averaged estimates of evolutionary rates for each morph. Unless otherwise indicated, all analyses were conducted using R v. 3.6.0 (R Core Team, 2019).

### Geophylogeny analyses

We could not carry out a formal biogeographical analysis, given that their high endemism and the complex pattern of underground connections between caves limits the establishment of reasonable biogeographical areas. However, we assessed the geographical structuring of *Sinocyclocheilus* diversification by building a geophylogeny on GenGIS v. 2.5.3 (Parks et al., 2013) based on the maximum credibility tree.

## RESULTS

The maximum credibility tree of *Sinocyclocheilus* is shown in Fig. 2 together with the reconstruction of ancestral states for eye-related morphs; we consider four major clades (A,B,C,D), as previously reported by other authors (Zhao and Zhang, 2009). Despite the inherent uncertainty in ancestral state reconstructions, it is clear that Blind species evolved at least three times in *Sinocyclocheilus*. Two of these events involved single species evolving from Normal-eyed ancestors, namely *S. xunlensis* and *S. anophthalmus*. On the other hand, the third lineage of blind *Sinocyclocheilus*, Clade B, includes several closely related species of Blind, Micro-eyed and a few Normal-eyed species, with two cases of reversal from either Micro-eyed or Blind to Normal-eyed morphs, namely *S. zhenfengensis* and *S. brevibarbatus* (Fig. 2). All four clades contain Regressed-eyed species and comparatively, clade Blind contains the most cases. Interestingly, clade B originated around the time of the beginning of the aridification process in China in the late Pliocene, whereas the other two transitions to blind species were much more recent (Fig. 2).

**Fig. 2.**
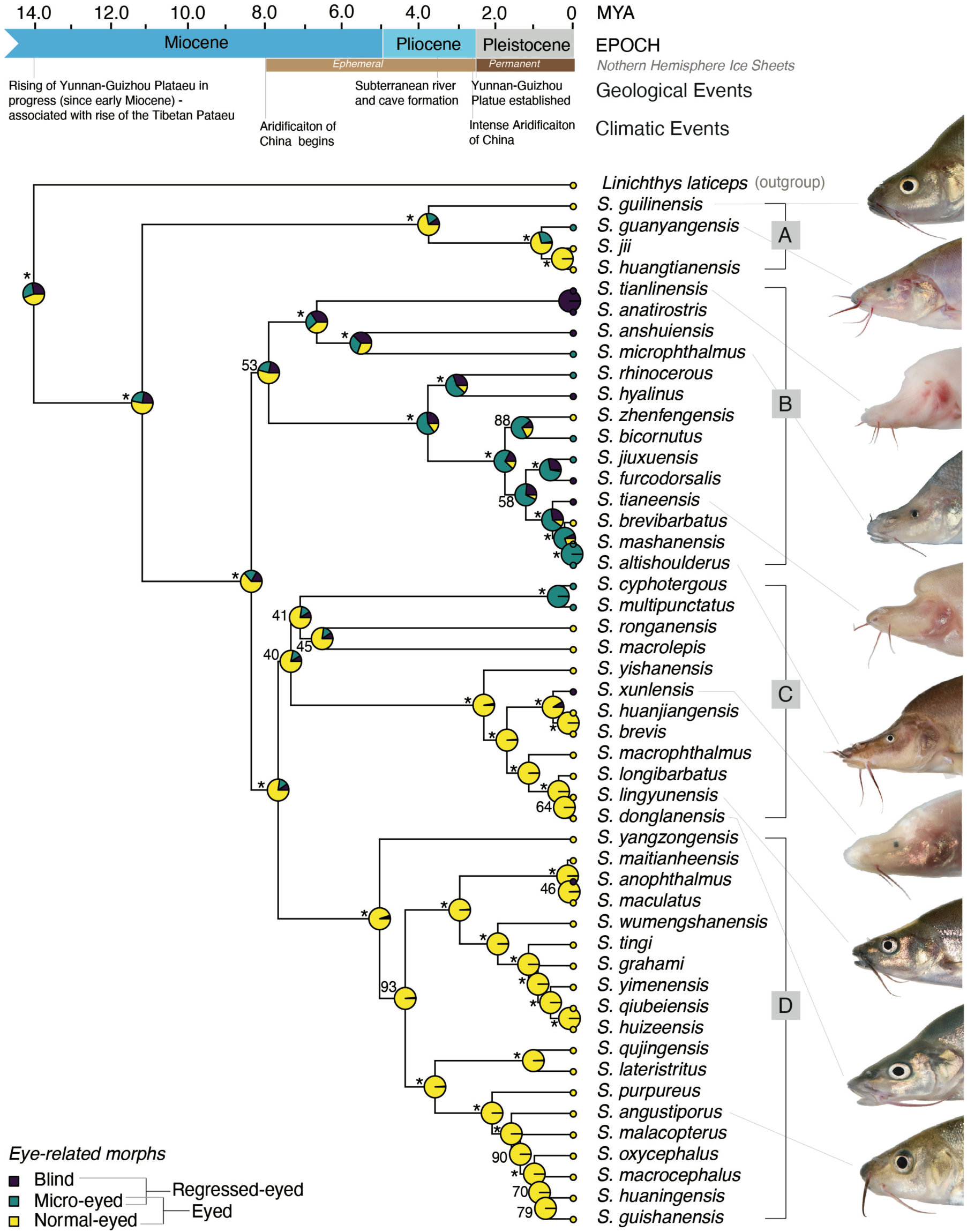
Ancestral state reconstruction of morphology (Blind, Mico-eyed & Normal-eyed) on a time calibrated phylogeny. Posterior probabilities of node support also shown on the tree. **Maximum-likelihood reconstructions for the ancestral state of the eye-trait morphology (Blind, Micro and Normal-eyed morphs) on a time calibrated phylogeny**. A, B, C and D are four major clades. Posterior probabilities of node support values of 100% are indicated by *. Key events of *Sinocyclocheilus* evolution includes; at least three independent evolutionary events for Blind morphs; Blind, Micro-eyed and a few Normal-eyed species in clade B, with two cases of reversal from either Micro-eyed or Blind to Normal-eyed species.

Interestingly, the evolution of body size and eye diameter seem to have often involved reversals, with little correspondence between body size (Fig. 3A) or eye diameter (Fig. 3B) and their corresponding morphs, except for the case of blind species for which eye diameter is necessarily zero. On the other hand, morphs are clearly distinguished when eye diameter and body size are visualized simultaneously (Fig. 3C), which suggests that the evolution of different morphs is achieved by altering the relationships between body size and eye diameter. Habitat associations traced on the phylomorphospace (Fig. 3D) indicates that species with regressed eyes and small-to-medium body sizes are obligate cave dwellers. However, normal eyed species can be Troglodytic, Troglophilic or surface dwellers regardless of their body size. Interestingly, all horned species are obligate cave dwellers while all cave species are not horned (Fig. 3E).

**Fig. 3.**
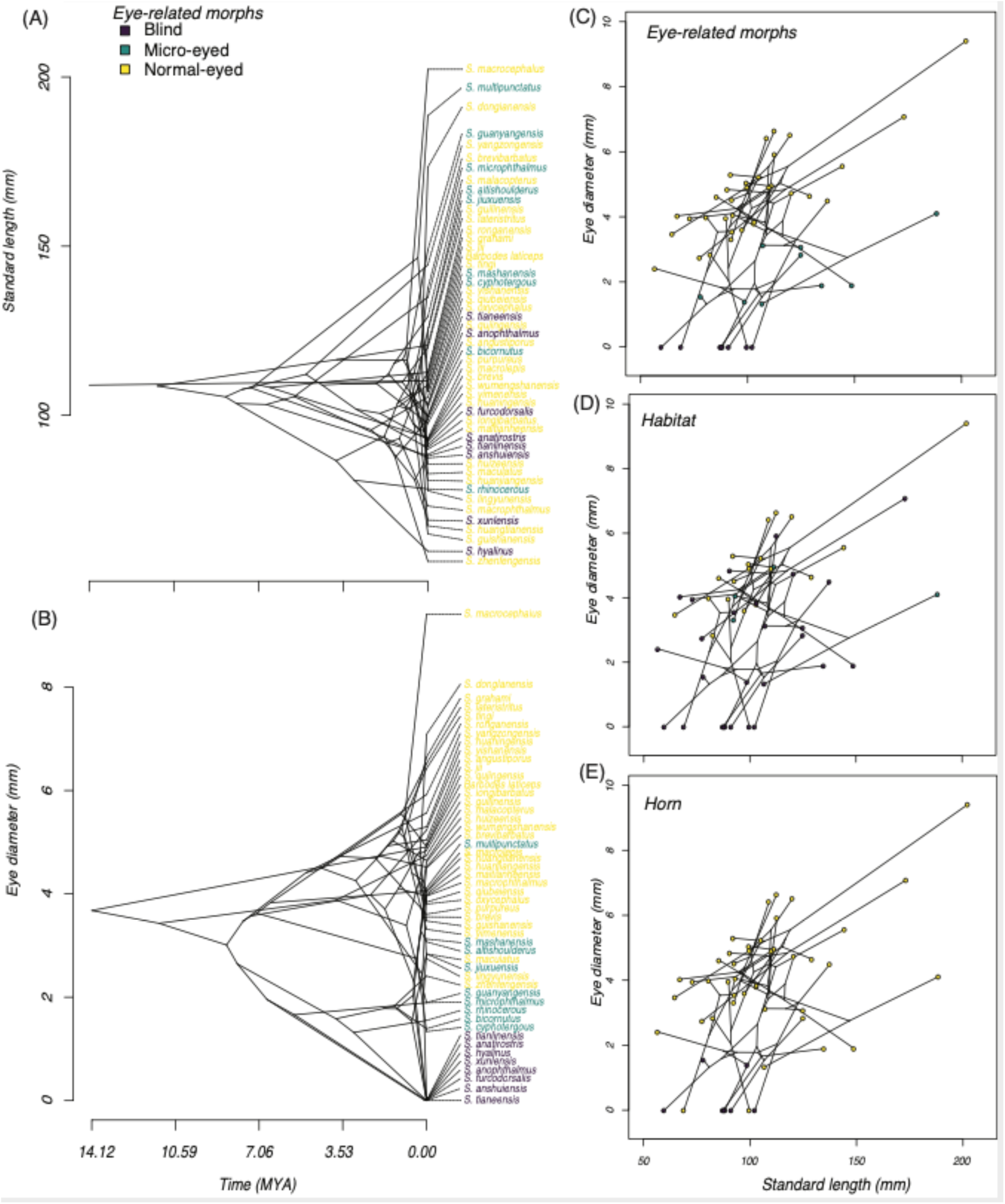
Temporal patterns of body size and eye diameter in *Sinocyclocheilus* from the perspective of their eye trait related morphs. The traitgrams suggests that the evolution of different morphs is attained by altering the allometric relationships between body size and eye diameter. A. Traitgram of body size; B. traitgram of eye size; C. Eye related morphs traced on the phylomorphospace indicating clear separation of the three morphs in the morphospace. (D)Habitat associations traced on the phylomorphospace showing species having eye diameter <3mm and small to medium body sizes are obligate cave dwellers whereas species with eye diameter >3mm can be Troglodytic, Troglophilic or Surface dwellers regardless of their body size (E) Horn related morphs traced on the phylomorphospace indicating the presence of a horn in smaller fish with reduced eye size. Horned species are Troglodytes.

A more precise description of the overall changes associated with different morphs can be visualized in the projections build from the geometric morphometrics analyses (Fig. 1B). The first PC, which accounted for approximately 32% of the variance in the dataset (see Table S3), tended to distinguish the slender Normal-eyed species on the left and Micro/Blind species on the right, which were characterized by changes in shape and widening of the anterior dorsal area between mouth and beginning of the dorsal fin of their body, resulting in a shift from the fusiform shape of the Normal-eyed forms to a more “boxy” form of the Micro-eyed and Blind forms. The second PC, which explained approximately 17% of the variance in the dataset, emphasized the differences in the type of dorsoventral broadening of the mid-section between morphs, with a shortening of the tail region (Fig. 1B). The variation in this axis is very high among the Micro-eyed and Blind forms when compared to the Normal morphs.

The multiple-rate model of evolution provided the best fit to the data for all three quantitative traits (ΔAIC=2.7, 10.7 and 7.9 respectively for ED, sED and SL; Table 1), indicating that the evolution of different *Sinocyclocheilus* morphs was associated with significant changes in their evolutionary rates. However, there were intriguing differences between traits in their rates (Table 2). The rates of evolution of eye diameter and standardized eye diameter were similar between Normal-eyed and Micro-Eyed species, but increased between 3.3 to 12.56 times during shifts towards Blind species.

**Table 1.**
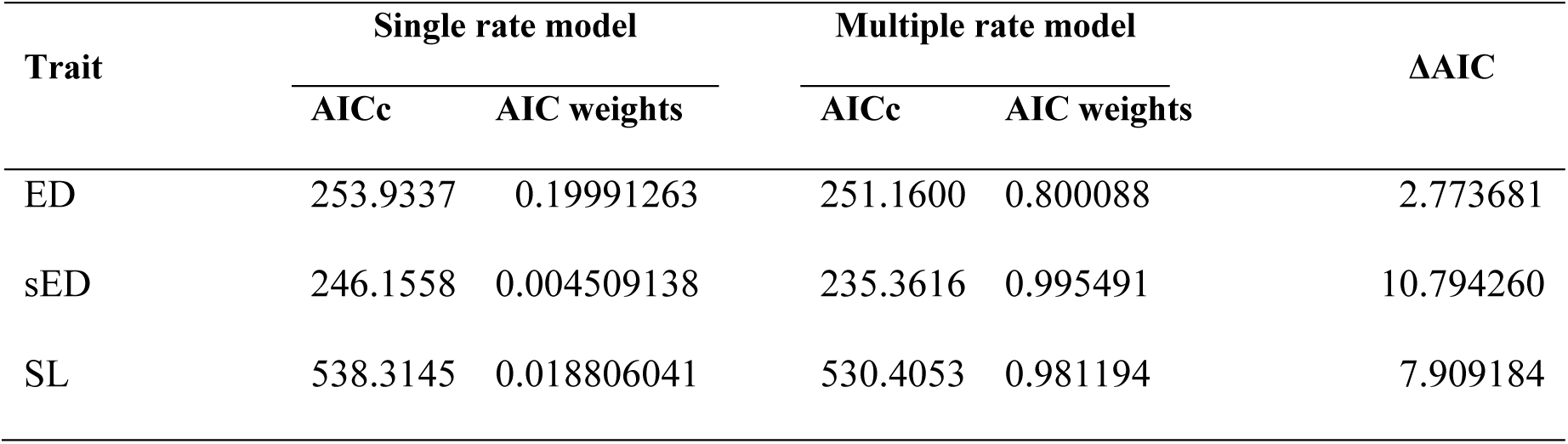
Model fit and estimated Brownian rate parameters for three traits (ED, sED and SL) in eye related morphs of *Sinocyclocheilus*. Multiple-rate models of evolution providing the best fit to the data for all three quantitative traits with ΔAIC >2.

**Table 2.** Model averaged rate parameters for the measured traits in eye related morphs of *Sinocyclocheilus*. Normal-eyed and Micro-Eyed species indicate similar evolutionary rates with marked shifts towards Blind species.

| Trait | Model Averaged Rate |  |  |
| --- | --- | --- | --- |
|  | Blind | Micro-eyed | Normal-eyed |
| SL | 45.3506 | 1597.9230 | 1275.5080 |
| ED | 11.3935 | 2.9984 | 3.8872 |
| sED | 24.9441 | 1.6925 | 2.2773 |

Geophylogeny represents the phylogeny overlaid across the geographic location of each species, where phylogenetic clustering is evident across the landscape. Considering the distribution of *Sinocyclocheilus*, we mainly see a pattern where the basal, Normal-eyed morphs are placed in the east, a substantial portion of Blind/Micro-eyed (Regressed-eyed) species are in the center, and Normal-eyed morphs are predominant towards the western mountains (Fig. 4).

**Fig. 4.**
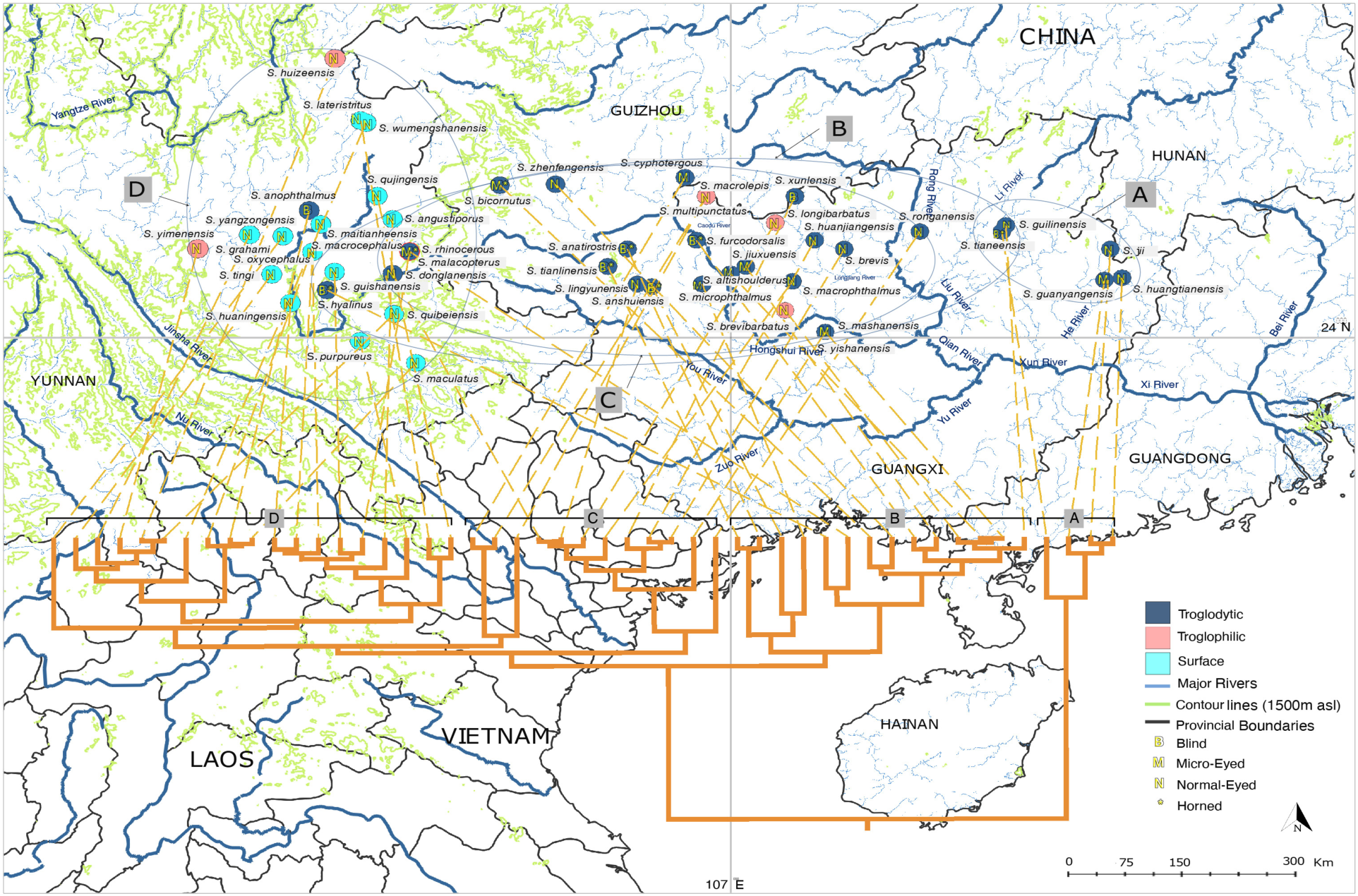
Geophylogeny, the phylogeny laid across the geographic distribution of the species considered in the analysis, with eye-related morphology, habitats, and horn-existence of the species traced. A pattern where basal, Normal-eyed, Troglodytic species are placed in the east, predominantly Blind/Micro-eyed/Normal-Eyed, Troglodytic species in the center and Normal-eyed, Surface dwelling species towards the western mountains is evident, indicating an East to west dispersion of the genus *Sinocyclocheilus* across South and South Western China. Eye specializations mostly occurred in Clade B, and horn evolution occurred exclusively in Clade B, within the Central range of the *Sinocyclocheilus* distribution.

## DISCUSSION

### Habitat utilization in context of eye-morphology

Integrating evolution of eye size and habitat manifests interesting and previously unrecognized evolutionary patterns in the evolution of *Sinocyclocheilus*. The Eye-size based ancestral reconstruction suggests the base of the phylogeny is most likely an Eyed species (i.e. Normal- or Micro-), but habitat reconstructions places, with high probability, Troglodytic species at the base (Fig. S1). This suggests an ancestral Eyed-species evolved a Troglodytic habit before they became blind. This may be an example of preadaptation in *Sinocyclocheilus*, i.e., the advancement of a functional change with little or no evolutionary modification (Ardila, 2016). In *Astyanax* cavefish, surface-dwelling forms are scotophilic, they prefer to remain away from direct light suggesting that scotophilia may be preadaptive for colonizing the dark, cave environment (Espinasa et al., 2001). In *Sinocyclocheilus*, since a basal (eyed) species demonstrated preference for the cave habitat, this preadaptation to darkness may hint towards why certain species tend to become cave-dwellers while others do not. This pattern is supported by two principal lines of evidence. First, most of the basal species are eyed, and Troglodytic (except for one species with an unusual eye-related polymorphic condition that we discuss below). Second, the most westward group (Clade D; Normal-eyed Surface fish), re-colonized caves whenever cave habitats were available within that area, suggesting a strong predisposition for cave-dwelling across all *Sinocyclocheilus*. In other words, when caves were present, members of *Sinocylocheilus*, irrespective of eye-related condition, preferred the cave habitat.

The preference for caves may not be a preference for darkness, but in fact a preference for depth, in search of water for survival. In a karstic environment where drying of surface running water is common, a preference for such deeper habitats may have provided an evolutionary advantage. In the presence of an array of subterranean waterways, such a predisposition would have given rise to the eye-regressed forms living close to, or associated with, caves that are characteristic of the genus.

Furthermore, apart from the Troglodytic and Troglophilic species of Clade D, some of the putative Surface species of Clade D are often observed at the entrances of caves or at windows to subterranean rivers (Zhao and Zhang, 2009). Hence, with more intensive ecological studies, some species recognized as Surface species may indeed be Troglophilic species, bolstering the notion that *Sinocyclocheilus* are predisposed to seek deeper waters of the karstic caves.

Resource utilization plays a key survival role in harsh environments (Culver and Pipan, 2009). Some of the Troglophilic, eyed-species are nocturnal, emerging from submerged caves, presumably to feed at night to reduce competition from other non-cave inhabiting fish species (personal observations). Some species like *S. altishoulderus, S. donglanensis* (Romero et al., 2009), *S. bamaensis* (Su et al., 2003), *S. malacopterus* (Chen et al., 2017) and *S. longibarbatus* (personal observation, video evidence as Supplementary information); are known to come out of caves during the high water season, presumably to feed and breed. This explains dependence on the cave as a diurnal refugium, from where these species can exploit the surface habitats at night. Strategies such as this, where multiple resources are utilized simultaneously, points to the adaptability of some *Sinocyclocheilus* species, resulting in their evolutionary success in a harsh and changing environment. Hence, the cave entrances are possibly an important ecotone that is important in *Sinocyclochelius* diversification and conservation.

Season and time of day seem to be important factors in determining habitat utilization patterns, but this level of resolution in habitat data is not currently available for a majority of the species to carry out a comprehensive habitat analysis across the diversification – indeed, many species are known only from one or a few specimens (Zhao and Zhang, 2009).

### Adaptations in the light of geophylogeny

In combination with the data analyzed, basal *Sinocyclocheilus* (Clade A) are Normal-Eyed, predominantly cave dwelling and non-horned species from the Eastern region of their distribution. As pointed out, this suggests that the earliest ancestors of *Sinocyclocheilus* species where Normal-eyed and but still lived in close association with caves. The ancestral reconstructions suggest that the affinity to caves would have evolved early and is present in most *Sinocyclocheilus*. The clade comprising basal species are from the east of the *Sinocyclocheilus* distribution, i.e. the He Jiang and Gui Jiang river basin in northeastern Guangxi, and hence, it seems that the diversification of these fish occurred from East to West (Fig. 4).

Within this predominantly Normal-eyed clade (Clade A), there is an exception, *Sinocyclocheilus guanyangensis*, a species that we coded as Micro-eyed, has Normal-Eyed, Micro-eyed and effectively Blind morphs within the same population – polymorphic for this trait. But these blind morphs have their eyes completely covered by skin and the Micro-eye is not itself degenerate. This kind of condition has been noted in several other taxa also (*S. xunlensis* and *S. flexuodorsalis –* not available for our analysis), but is uncommon. This suggests a degree of polymorphism for this trait, suggesting that the earliest ancestors of *Sinocyclocheilus* may have been able to lose or gain eyes relatively easily as an adaptation to local conditions, this ability appears several times within this cave-driven diversification.

The major adaptation for cave dwelling evolves predominantly in the expansive karstic area in northwestern Guangxi (associated with the Liu Jiang basin and Hongshui river basin joining the main Xijiang River system from the North), in Clade B, the southeastern corner of Guizhou province (upper reaches of Hongshui River) and the northeastern plateau of Yunnan province. This region can be considered the center for novel adaptations for *Sinocyclocheilus*, where these fishes express their full morphological diversity, blindness, micro-eyedness, and their remarkable horns. In the shape-related analyses, these species cluster on the right of morphospace (Fig. 1B). The deeper caves and extensive subterranean river system associated with the Guangxi plains (Zhao and Zhang, 2009) would have facilitated this extensive adaptive diversification (Fig. 4).

The karstic northwestern region that the Guangxi-dominated Clade (Clade B) experiences drought conditions during much of the year, and one of the major sources of rain for the region is through storms sweeping from the southeast that are strong enough to persist through the vast plains of Guangxi, mainly from April to August (Zhao and Zhan, 2009). During unfavorable periods, these fishes seem to have found refuge in the subterranean caves. The morphologically most diverse clade being present in the region where the climatic conditions are most unfavorable for surface fish, reinforces the notion that *Sinocyclocheilus* species adapted to life in caves as climatic refuges (Zhao and Zhang, 2009).

The distribution of Clade C, characterized by mostly Normal-eyed but Troglodytic species largely overlaps Clade B. In Clade C, the single Blind species (*S. xunlensis*) and the two Micro-Eyed species (*S. cyphotergous* and *S. multipunctatus*) are shown within a narrow geographic area on the Liu Jiang and Hongshui river system (Fig. 4).

Species in the clade that is found in the west (Clade D), predominantly on the hilly terrain of Yunnan plateau, are predominantly Normal-eyed Surface species lacking horns (Fig. 1, Fig. 4). However, wherever there are cave habitats and subterranean river systems, some of these putative surface species have become facultative or obligate cave species. The obligate cave species found within the region, *S. anophthalmus*, is blind. However, this Blind species stands clustered with the Normal-eyed morphs in the morphospace, signifying that the shape of the species has not extensively changed, possibly due to recent (Pleistocene) invasion of the cave habitat from a Normal-eyed ancestor (Fig. 2) – time since becoming blind is not been long enough for change into the box-like shape of the Blind species of Clade B.

Horn distributions show several peculiar trends. In most *Sinocyclocheilus* species, a prominent hump is present (He et al., 2013). However, this hump is markedly low in the Normal-eyed surface inhabiting species of the Yunnan clade (Fig. 4). For the species that bears a horn, the horn represents the region in which the dorso-frontal hump is present, and always occurs before the hump begins, at the boundary of the edge of the dorsal skull. The exception to this is *S. cyphotergous* (Clade C), where a horn like structure is placed on top of the hump. *S. cyphotergous*, a species found in clade C, is phylogenetically separate from other horned species, suggesting that the origins of the “horn” for this species is evolutionarily different from the other horned species (Fig. S2). Though the function of the horn remains unknown (protection of head, anchoring in strong current and protection of head has been suggested; Zhao and Zhang, 2009), functionally this structure may be similar to other species, if it is actually anchoring in strong current is the main function.

The rates of evolution of various traits show some incongruent (non-allometric), but interesting patterns that can be explained in the context to adaptations to a Troglodytic condition. The rates of evolution in eye diameter are similar between Normal- and Micro-Eyed species but, increases dramatically (3.3-12.56 times) with shifts towards blind forms. However, body size evolution for these morpho groups shows a reversed pattern, with a 0.03 decrease in body size evolution in the Blind morphs compared to the eyed-morphs. These patterns in rate variation suggest that the evolution of Blind morphs to a Troglodytic habitat were simultaneously associated with an increase in the rate of evolution of the eye-size degeneration and a decrease in the rate of body size evolution. The smaller body size resulting from a sluggish rate of change will facilitate both navigation within constricted spaces and sustenance on a limited supply of resources expected to be experienced in subterranean habitats.

Much of our collective knowledge of the patterns and mechanisms of regressive evolution come from studies of animals that have colonized the subterranean biome. Within this group, a several studies have focused on the Mexican tetra, *Astyanax mexicanus* (Jeffery, 2009). This natural animal model system comprises multiple cave-adapted morphs and a surface-dwelling morph that resides in near the caves themselves (Gross, 2012). Since the discovery of *Astyanax* cavefish in 1936, countless studies have provided insight to the developmental and genetic bases for cave-associated traits (Hubbs and Innes, 1936). Indeed, much of this insight has emerged from the interbreeding studies of conspecific cave and surface morphs (reviewed in Wilkens, 2016). However, several aspects of regressive evolution and troglomorphic adaptation remain unresolved. Owing to several of the differences with *Astyanax*, we argue that *Sinocyclocheilus* is well-positioned to provide important new insights to broader patterns of diversification and adaptation in cave-dwelling organisms.

For instance, *Astyanax* represents a single species with 30 different interfertile populations from a relatively small geographic location (Wilkens and Strecker, 2017). In contrast, *Sinocyclochelius* harbors about 75 species (49 in our study) inhabiting diverse geographic biomes across a much larger geographic area. Moreover, while *Astyanax* cavefish converge on similar phenotypes (regressed vision and pigmentation), they are not regarded as having numerous morphological novelties. In contrast, *Sinocyclocheilus* species have recurrently evolved a unique “horn” (Soares et al., 2019) from several eyeless species. Similarly, the larger number of *Sinocyclocheilus* species allows keener resolution for understanding broad phylogenetic processes, such as trait reversals and directions of diversification. Although a reversal from an eyeless to an eyed form has been reported for one cave population in *Astyanax* (Caballo Moro; Krishnan and Rohner, 2017), this phenomenon appears to be much less common than in *Sinocyclocheilus*. Additionally, a clear polarity of diversification is lacking in *Astyanax* cavefish, rather ancient stocks of surface-dwelling forms appear to have recurrently invaded caves to the east (i.e., Sierra de El Abra caves), with more recent invasions having occurred in the northern (Sierra de Colmena) and the western caves (Sierra de Guatemala; Bradic et al., 2012). However, the well-characterized gene flow between the cave and surface waters obscures the ability to understand clear boundaries between different cave groups. Further, most *Astyanax* cave populations are believed to have diverged over the course of the last ∼200 – 500 Ky (Herman et al., 2018). By contrast, *Sinocyclocheilus* species are much older, and therefore one can determine how longer-term processes unfold in these cave-dwelling animals. Thus, despite clear phylogenetic differences between *Astyanax* and *Sinocyclocheilus*, both genera have the ability to provide complementary and critical insights to the processes underlying cave evolution and diversification.

The integration of morphology, phylogeny, rate analyses, dating and distribution show not only several remarkable patterns of evolution, but also interesting exceptions to these patterns that signifies the diversification of *Sinocyclocheilus* as a unique model system to study evolutionary novelty.

## ACKNOWLEDGEMENTS

We thank the following institutions and individuals: funding from Guangxi University Special Talent Recruitment Grant to MM; funding from National Natural Science Foundation of China (31860600 and Guangxi Natural Science Foundation (2017GXNSFFA198010) JY; Shipeng Zhou, Bing Chen, Dan Sun, Jayampathi Herath and Amrapali Rajput for assistance in the field; Ethical review approval from Guangxi University.

## Supplementary Information

**Fig. S1.**
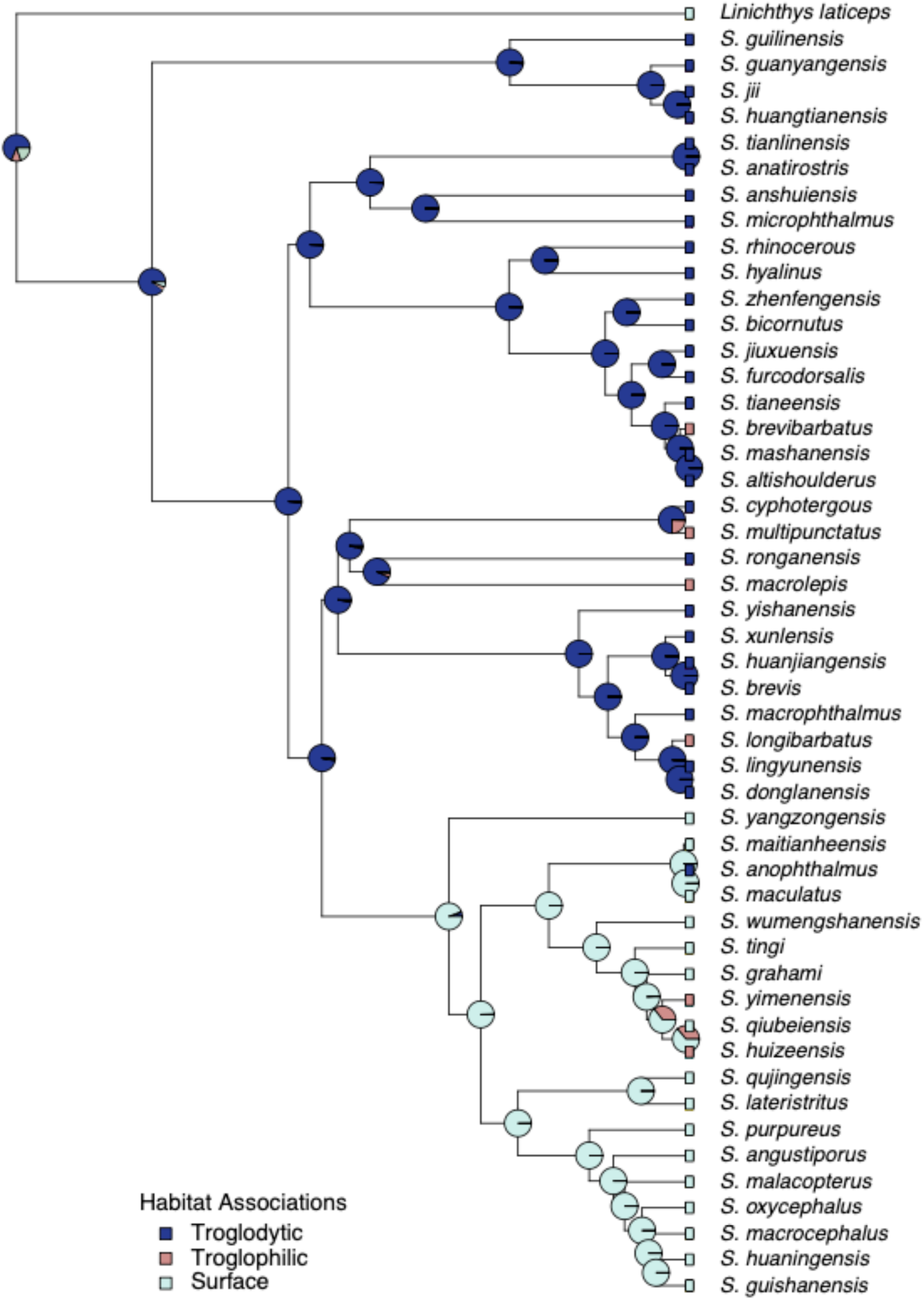
Maximum-likelihood reconstructions for the ancestral states of the habitat occupation (Troglodytic, Troglophilic and Surface) on the phylogeny of the genus *Sinocyclocheilus*.

**Fig. S2.**
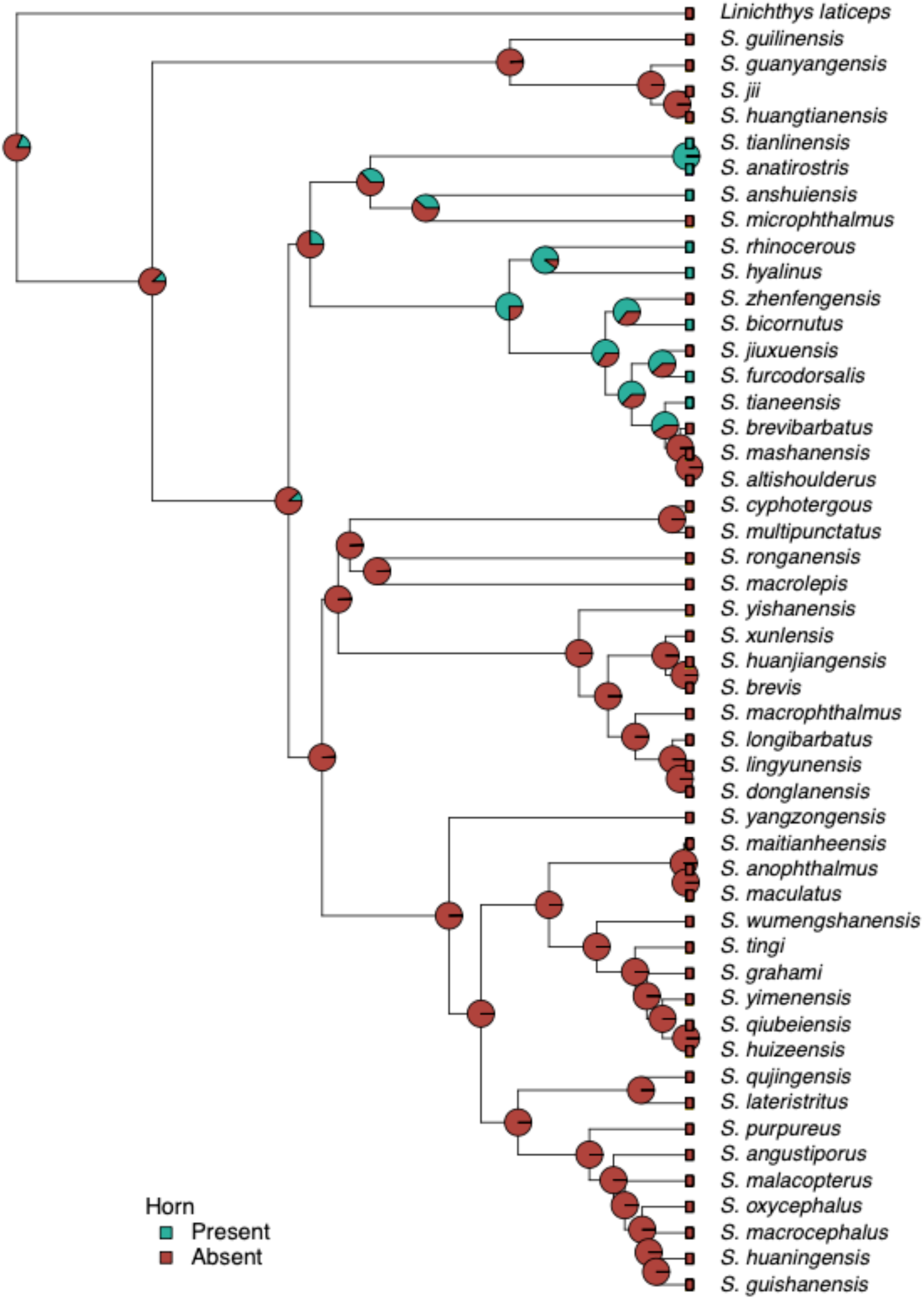
Maximum-likelihood reconstructions for the ancestral state of the horn related trait (presence/absence of horn) on the phylogeny of the genus *Sinocyclocheilus*.

**Table S1.**
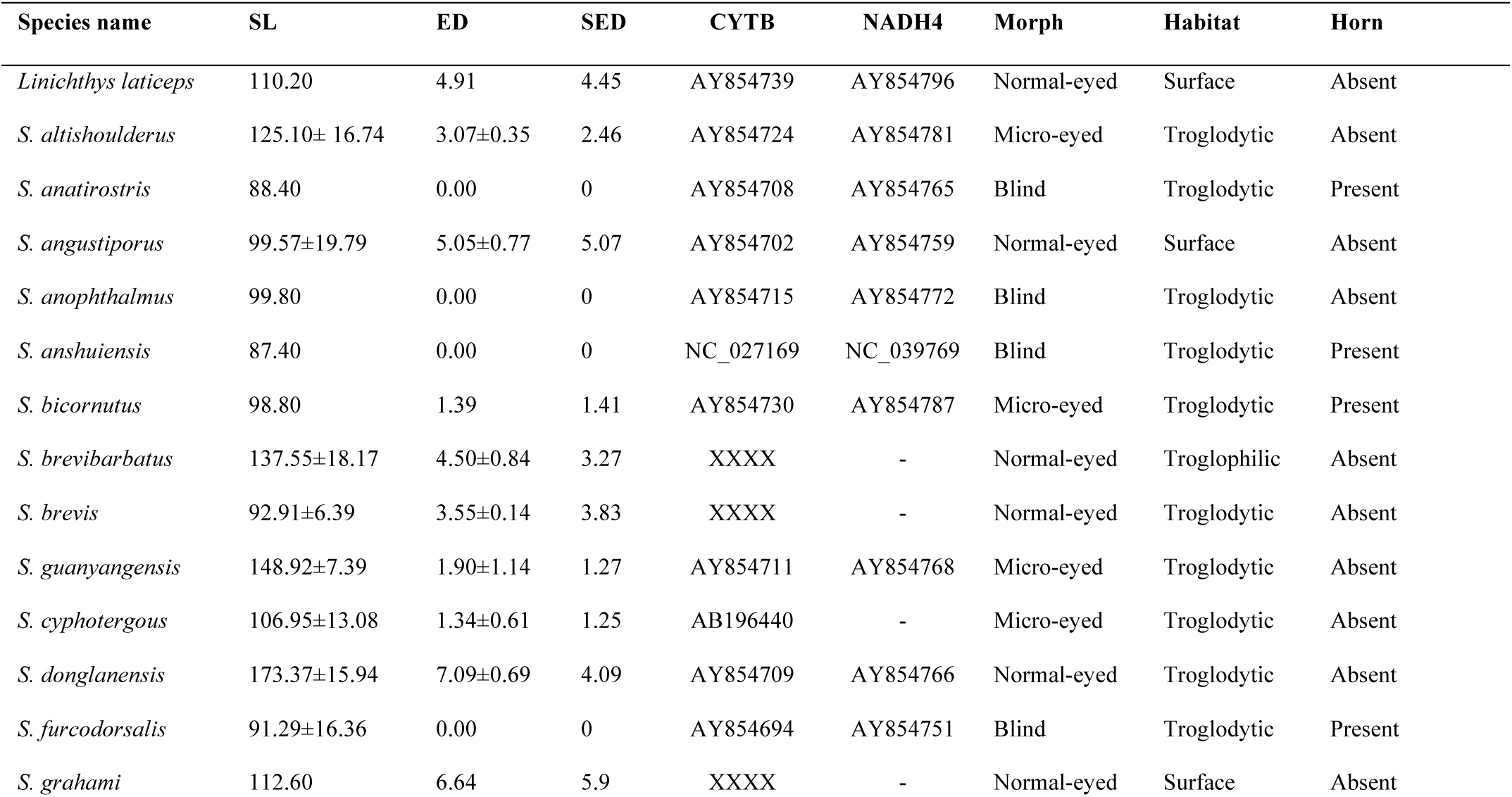
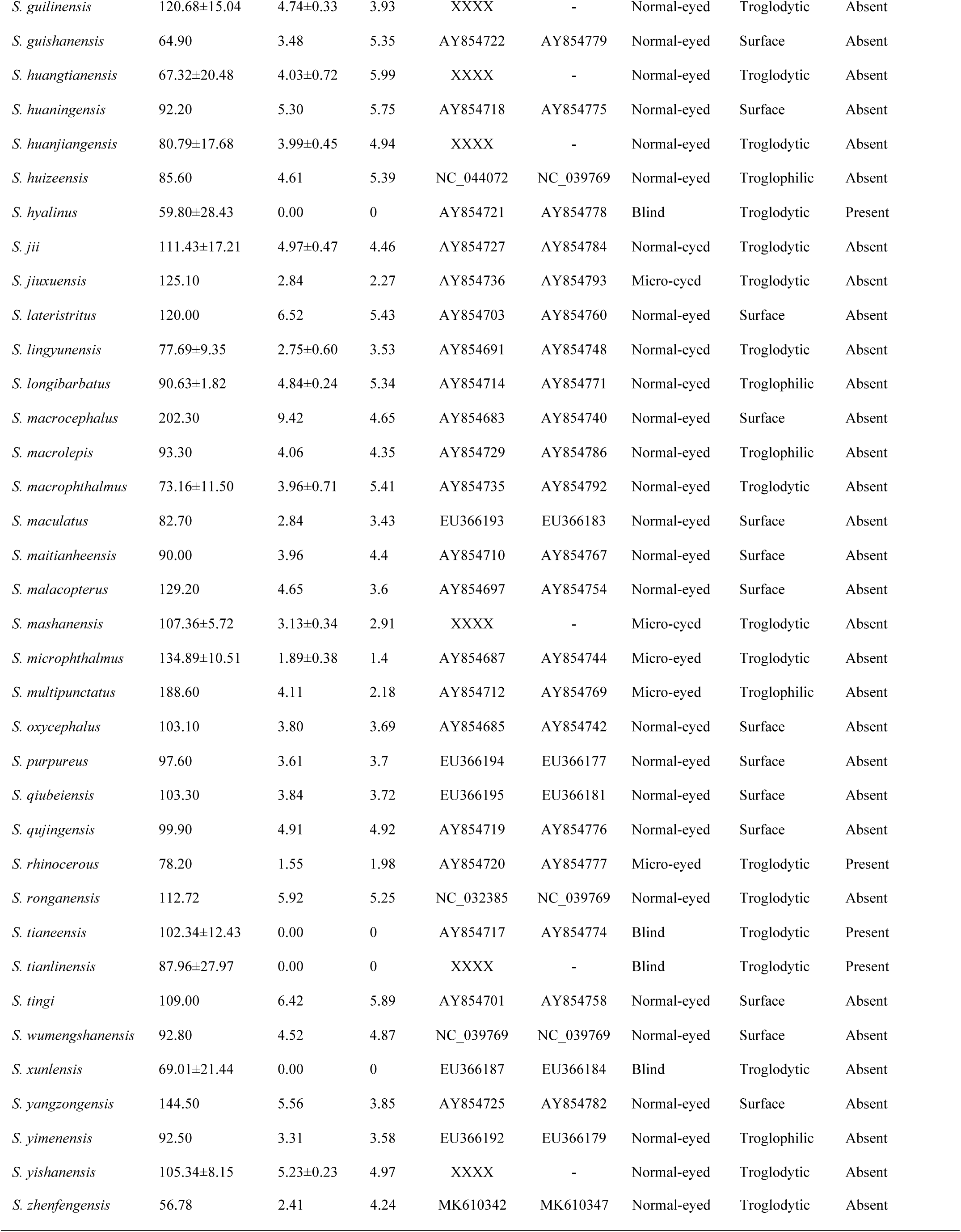
Species information and GenBank accession numbers of two mtDNA fragments (*NADH4* and *cytb*) of 39 *Sinocyclocheilus* species. The information of the outgroup species *Linichthys laticeps* (Cyprinidae) is also indicated along with calculated standard length (SL), eye diameter (ED), standard eye diameter (sED), discrete trait categories related to eye morphology (Blind, Micro-eyed, Normal eyed), presence or absence of a horn and habitat occupation (Troglodytic, Troglophilic and surface) for all species used in the analysis. Accession numbers indicated as XXXX will be accessible upon the acceptance of the manuscript.

**Table S2.** Information of digitized images used in the morphometric geometric analysis. Table indicates the voucher number of the specimen used for the analysis and the reference from which the image was obtained. Images photographed during the current study are also indicated with voucher numbers stated as GXUXXX (GXU: Guangxi University, China).

| Species name | Reference | Voucher # |
| --- | --- | --- |
| *Linichthys laticeps* | Zhang, E., and F. Fang. 2005. *Linichthys*: A New Genus of Chinese Cyprinid Fishes (Teleostei: Cypriniformes). *Copeia* 2005:61–67. | *IHB 78X6242* |
| *S. altishoulderus* | Guangxi University (this study). | *GXU001, GXU002, GXU003* |
| *S. anatirostris* | Romero, A., Y. Zhao, and X. Chen. 2009. The Hypogean fishes of China. *Environ Biol Fish* 86:211–278. | *IHB 84VII255,* |
| *S. angustiporus* | Guangxi University (this study). | *GXU007, GXU008 , GXU009* |
| *S. anophthalmus* | Romero, A., Y. Zhao, and X. Chen. 2009. The Hypogean fishes of China. *Environ Biol Fish* 86:211–278. | *KIZ 865949* |
| *S. anshuiensis* | Xi, G., W. Tie-Jun, W. Mu-Lan, and Y. Jian. 2013. A new blind barbine species, *Sinocyclocheilus anshuiensis* sp. nov.(Cypriniformes: Cyprinidae) from Guangxi, China. Kunming Institute of Zoology, Chinese Academy of Sciences. | *12070276* |
| *S. bicornutus* | Romero, A., Y. Zhao, and X. Chen. 2009. The Hypogean fishes of China. *Environ Biol Fish* 86:211–278. | *IHB 12209043-9o5o241* |
| *S. brevibarbatus* | Guangxi University (this study). | *GXU010, GXU011, GXU012* |
| *S. brevis* | Guangxi University (this study).<br>Romero, A., Y. Zhao, and X. Chen. 2009. The Hypogean fishes of China. *Environ Biol Fish* 86:211–278. | *GXU013, GXU014*<br>*IHB12209033-* |
|  |  | 87087496 |
| *S. cyphotergous* | Romero, A., Y. Zhao, and X. Chen. 2009. The Hypogean fishes of China. *Environ Biol Fish* 86:211–278.<br>Huang, J., A. Gluesenkamp, D. Fenolio, Z. Wu, and Y. Zhao. 2017. Neotype designation and redescription of *Sinocyclocheilus cyphotergous* (Dai) 1988, a rare and bizarre cavefish species distributed in China (Cypriniformes: Cyprinidae). *Environ Biol Fish* 100:1483–1488. | IHB 12209040<br>ASIZB 204678 |
| *S. donglanensis* | Guangxi University (this study). | *GXU015, GXU016, GXU017* |
| *S. furcodorsalis* | Guangxi University (this study). | *GXU018, GXU019, GXU020* |
| *S. grahami* | Romero, A., Y. Zhao, and X. Chen. 2009. The Hypogean fishes of China. *Environ Biol Fish* 86:211–278. | *ASIZB 03496* |
| *S. guanyangensis* | Guangxi University (this study). | *GXU004, GXU005, GXU006* |
| *S. guilinensis* | Guangxi University (this study). | *GXU021, GXU022, GXU023* |
| *S. guishanensis* | Romero, A., Y. Zhao, and X. Chen. 2009. The Hypogean fishes of China. *Environ Biol Fish* 86:211–278. | *Li980514005* |
| *S. huangtianensis* | Guangxi University (this study). | *GXU024, GXU025, GXU026* |
| *S. huaningensis* | Romero, A., Y. Zhao, and X. Chen. 2009. The Hypogean fishes of China. *Environ Biol Fish* 86:211–278. | *ASIZB79228* |
| *S. huanjiangensis* | Guangxi University (this study). | *GXU027, GXU028, GXU029* |
| *S. huizeensis* | Cheng, C., P. Xiao-Fu, C. Xiaoyong, J. Li, L. Ma, and J. Yang. 2015. A new species of the genus *Sinocyclocheilus* (Teleostei: Cypriniformes), from Jinshajiang Drainage, Yunnan, China. *Cave Research* 1:1–4. | *KIZ2013001246* |
| *S. hyalinus* | Romero, A., Y. Zhao, and X. Chen. 2009. The Hypogean fishes of China. *Environ Biol Fish* 86:211–278. | *KIZ916001*<br>*Photograph in life* |
|  | You, H., C. Xiaoyong, X. Ti-Qao, and Y. Jun-Xing. 2013. Three-dimensional morphology of the *Sinocyclocheilus hyalinus* (Cypriniformes : Cyprinidae) horn based on synchrotron X-ray microtomography. Zoological research 34:128–134. |  |
| *S. jii* | Romero, A., Y. Zhao, and X. Chen. 2009. The Hypogean fishes of China. Environ Biol Fish 86:211–278.<br>Baradi, W., Z. Yahui, Z. Chunguang, D. Rongji, and A. Abdul. 2013. Anatomical Studies of the Olfactory Epithelium of Two Cave Fishes *Sinocyclocheilus jii* and *S. furcodorsalis* (Cypriniformes: Cyprinidae) from China. Pakistan Journal of Zoology 45:1091 – 1101. | *ASIZB62726*<br>*Photograph in life* |
| *S. jiuxuensis* | Romero, A., Y. Zhao, and X. Chen. 2009. The Hypogean fishes of China. Environ Biol Fish 86:211–278. | *ASIZB102260* |
| *S. lateristritus* | Romero, A., Y. Zhao, and X. Chen. 2009. The Hypogean fishes of China. Environ Biol Fish 86:211–278. | *IHB12209036-865027* |
| *S. lingyunensis* | Guangxi University (this study).<br>Romero, A., Y. Zhao, and X. Chen. 2009. The Hypogean fishes of China. Environ Biol Fish 86:211–278. | *GXU030*<br>*ASIZB 73038* |
| *S. longibarbatus* | Guangxi University (this study). | *GXU031, GXU032, GXU033* |
| *S. macrocephalus* | Romero, A., Y. Zhao, and X. Chen. 2009. The Hypogean fishes of China. Environ Biol Fish 86:211–278. | *IHB12209012-662001* |
| *S. macrolepis* | Romero, A., Y. Zhao, and X. Chen. 2009. The Hypogean fishes of China. Environ Biol Fish 86:211–278. | *IHB12209035-87IV457* |
| *S. macrophthalmus* | Guangxi University (this study). | *GXU034, GXU035, GXU036* |
| *S. maculatus* | Romero, A., Y. Zhao, and X. Chen. 2009. The Hypogean fishes of China. Environ Biol Fish 86:211–278. | *Li870808001* |
| *S. maitianheensis* | Romero, A., Y. Zhao, and X. Chen. 2009. The Hypogean fishes of China. Environ Biol Fish 86:211–278. | IHB12209039-874001 |
| *S. malacopterus* | Romero, A., Y. Zhao, and X. Chen. 2009. The Hypogean fishes of China. Environ Biol Fish 86:211–278. | KIZ775831 |
| *S. mashanensis* | Guangxi University (this study). | GXU037, GXU038, GXU039 |
| *S. microphthalmus* | Guangxi University (this study). | GXU040, GXU041, GXU042 |
| *S. multipunctatus* | Romero, A., Y. Zhao, and X. Chen. 2009. The Hypogean fishes of China. Environ Biol Fish 86:211–278. | ASIZB73000 |
| *S. oxycephalus* | Romero, A., Y. Zhao, and X. Chen. 2009. The Hypogean fishes of China. Environ Biol Fish 86:211–278. | IHB12209013-652047 |
| *S. purpureus* | Romero, A., Y. Zhao, and X. Chen. 2009. The Hypogean fishes of China. Environ Biol Fish 86:211–278. | IHB12209015-731004 |
| *S. qiubeiensis* | Romero, A., Y. Zhao, and X. Chen. 2009. The Hypogean fishes of China. Environ Biol Fish 86:211–278. | Li990527002 |
| *S. qujingensis* | Romero, A., Y. Zhao, and X. Chen. 2009. The Hypogean fishes of China. Environ Biol Fish 86:211–278. | ASIZB78790 |
| *S. rhinoceros* | Romero, A., Y. Zhao, and X. Chen. 2009. The Hypogean fishes of China. Environ Biol Fish 86:211–278. | ASIZB93907 |
| *S. ronganensis* | FuGuang, L., H. Jie, L. Xia, L. Tong, and W. YanHong. 2016. *Sinocyclocheilus ronganensis* Luo, Huang et Wen sp. nov., a new species belonging to *Sinocyclocheilus* Fang from Guangxi (Cypriniformes: Cyprinidae). Guangxi Academy of Agricultural Sciences 47:650–655. | 20151114001 |
| *S. tianeensis* | Guangxi University (this study). | GXU043, GXU044, GXU045 |
| *S. tianlinensis* | Guangxi University (this study). | GXU052, GXU053, GXU054 |
| *S. tingi* | Romero, A., Y. Zhao, and X. Chen. 2009. The Hypogean fishes of China. Environ Biol Fish 86:211–278. | ASIZB60227 |
| *S. wumengshanensis* | Romero, A., Y. Zhao, and X. Chen. 2009. The Hypogean fishes of China. Environ Biol Fish 86:211–278. | *KIZ82100006* |
| *S. xunlensis* | Guangxi University (this study). | *GXU046, GXU047, GXU048* |
| *S. yangzongensis* | Romero, A., Y. Zhao, and X. Chen. 2009. The Hypogean fishes of China. Environ Biol Fish 86:211–278. | *KIZ6351069* |
| *S. yimenensis* | Romero, A., Y. Zhao, and X. Chen. 2009. The Hypogean fishes of China. Environ Biol Fish 86:211–278. | *Li030509009* |
| *S. yishanensis* | Romero, A., Y. Zhao, and X. Chen. 2009. The Hypogean fishes of China. Environ Biol Fish 86:211–278. | *GXU049, GXU050, GXU051* |
| *S. zhenfengensis* | Liu, T., H. Q. Deng, L. Ma, N. Xiao, and J. Zhou. 2018. *Sinocyclocheilus zhenfengensis*, a new cyprinid species (Pisces: Teleostei) from Guizhou Province, Southwest China. J Appl Ichthyol 34:945–953. | *GZNU20120701001* |

**Table S3.**
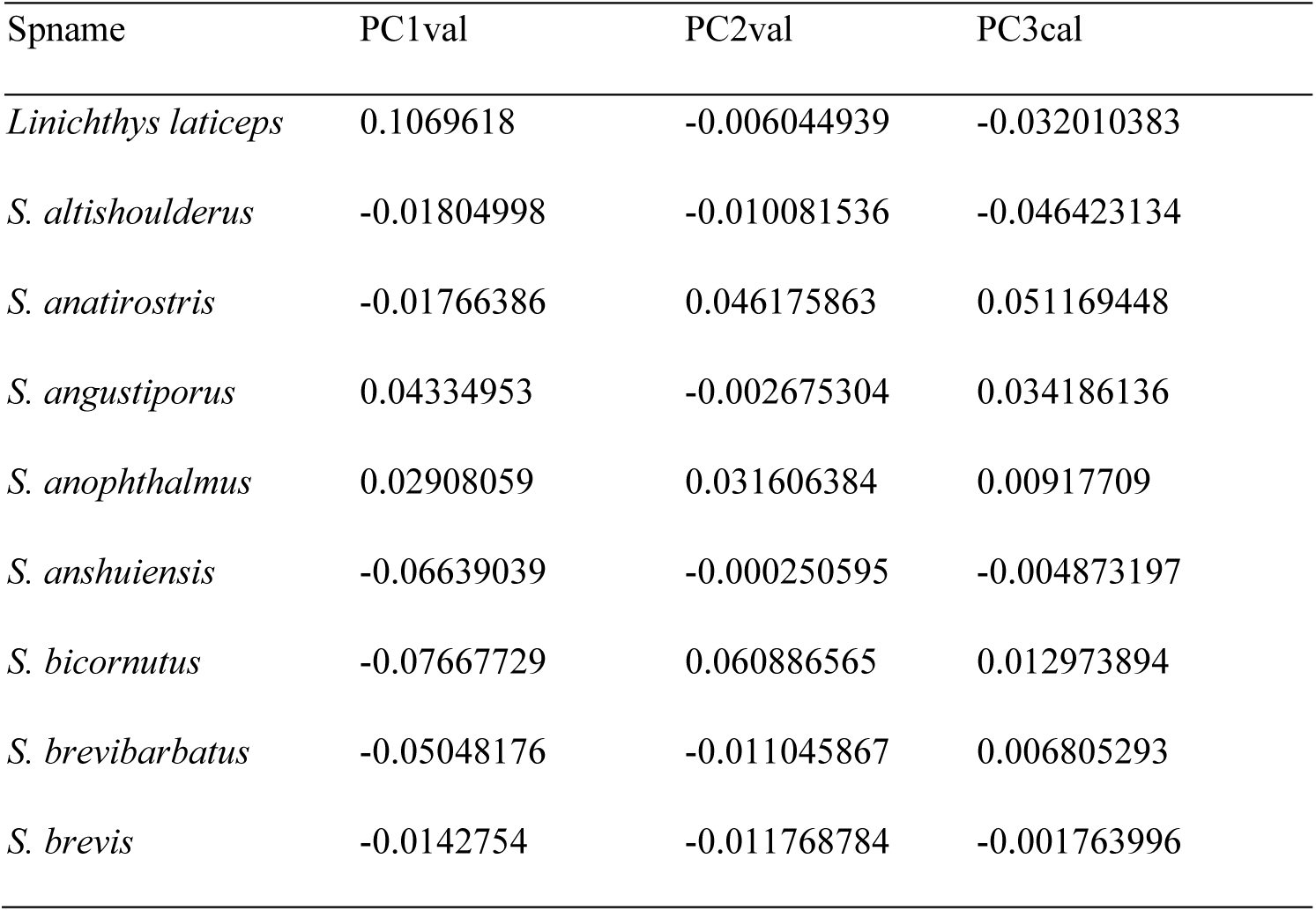
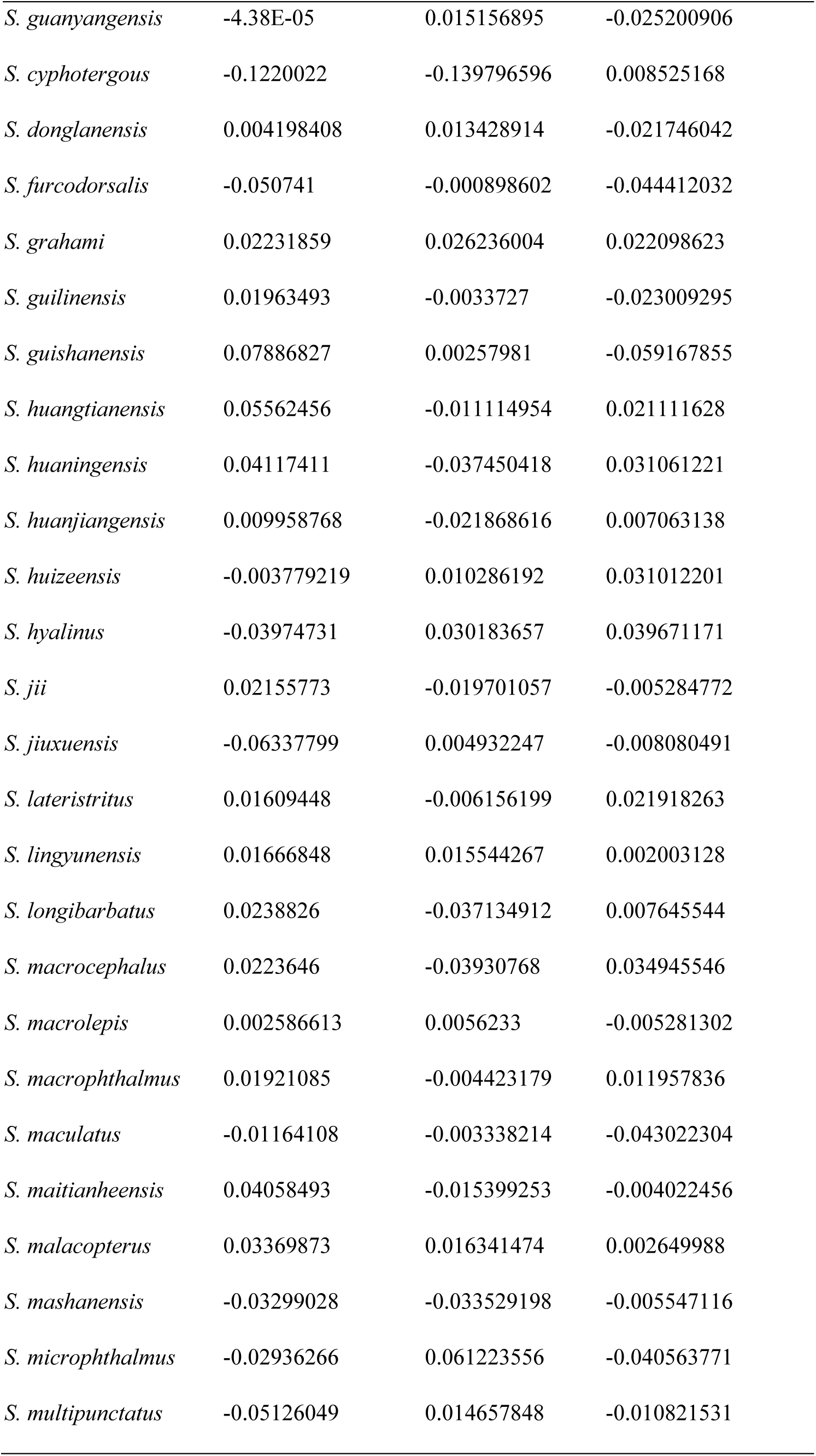
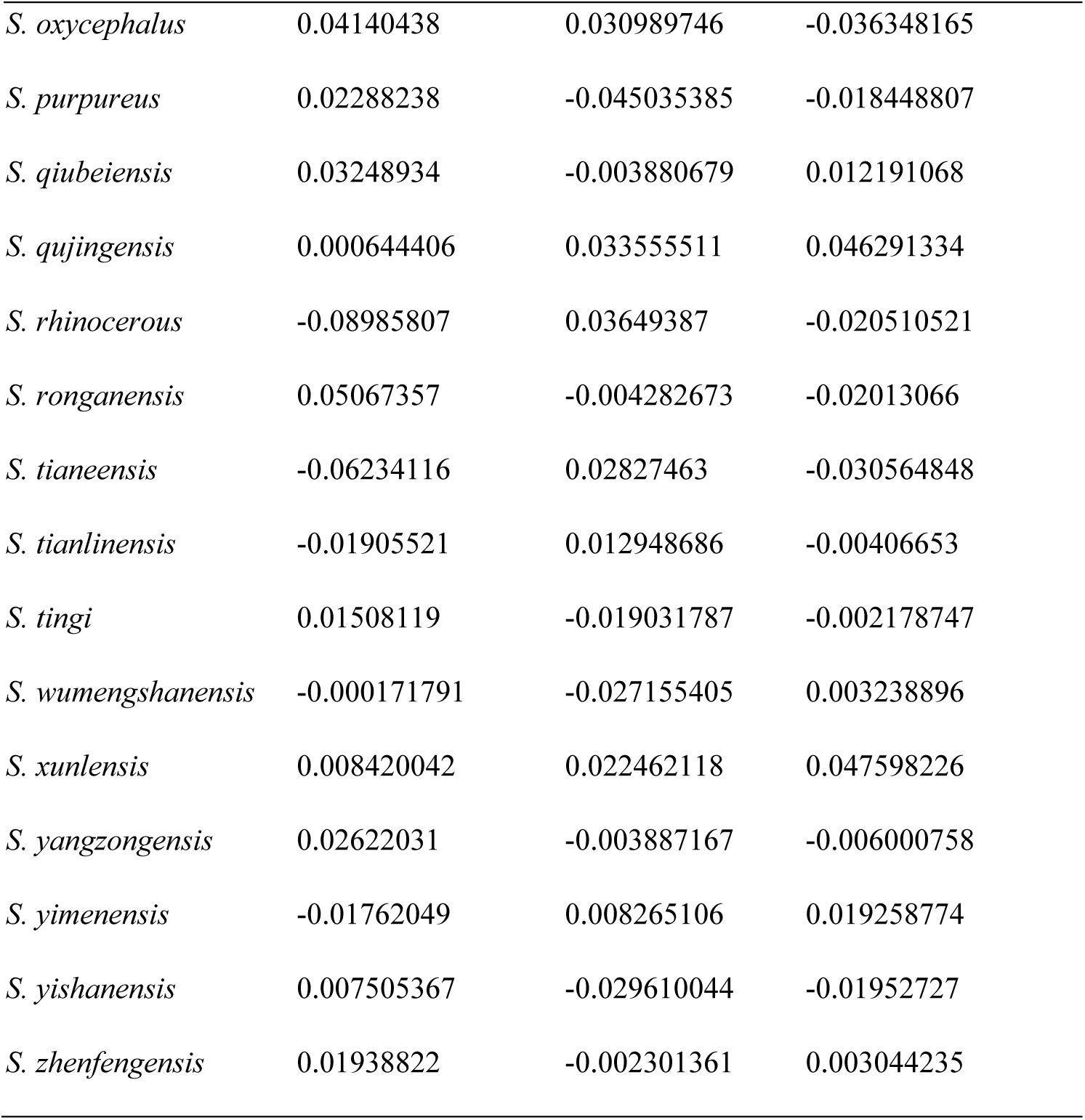
Calculated Principal Component values (PC1, PC2 and PC3) of all the specimens used in the current analysis

| Spname | PC1val | PC2val | PC3cal |
| --- | --- | --- | --- |
| *Linichthys laticeps* | 0.1069618 | -0.006044939 | -0.032010383 |
| *S. altishoulderus* | -0.01804998 | -0.010081536 | -0.046423134 |
| *S. anatirostris* | -0.01766386 | 0.046175863 | 0.051169448 |
| *S. angustiporus* | 0.04334953 | -0.002675304 | 0.034186136 |
| *S. anophthalmus* | 0.02908059 | 0.031606384 | 0.00917709 |
| *S. anshuiensis* | -0.06639039 | -0.000250595 | -0.004873197 |
| *S. bicornutus* | -0.07667729 | 0.060886565 | 0.012973894 |
| *S. brevibarbatus* | -0.05048176 | -0.011045867 | 0.006805293 |
| *S. brevis* | -0.0142754 | -0.011768784 | -0.001763996 |
| *S. guanyangensis* | -4.38E-05 | 0.015156895 | -0.025200906 |
| *S. cyphotergous* | -0.1220022 | -0.139796596 | 0.008525168 |
| *S. donglanensis* | 0.004198408 | 0.013428914 | -0.021746042 |
| *S. furcodorsalis* | -0.050741 | -0.000898602 | -0.044412032 |
| *S. grahami* | 0.02231859 | 0.026236004 | 0.022098623 |
| *S. guilinensis* | 0.01963493 | -0.0033727 | -0.023009295 |
| *S. guishanensis* | 0.07886827 | 0.00257981 | -0.059167855 |
| *S. huangtianensis* | 0.05562456 | -0.011114954 | 0.021111628 |
| *S. huaningensis* | 0.04117411 | -0.037450418 | 0.031061221 |
| *S. huanjiangensis* | 0.009958768 | -0.021868616 | 0.007063138 |
| *S. huizeensis* | -0.003779219 | 0.010286192 | 0.031012201 |
| *S. hyalinus* | -0.03974731 | 0.030183657 | 0.039671171 |
| *S. jii* | 0.02155773 | -0.019701057 | -0.005284772 |
| *S. jiuxuensis* | -0.06337799 | 0.004932247 | -0.008080491 |
| *S. lateristritus* | 0.01609448 | -0.006156199 | 0.021918263 |
| *S. lingyunensis* | 0.01666848 | 0.015544267 | 0.002003128 |
| *S. longibarbatus* | 0.0238826 | -0.037134912 | 0.007645544 |
| *S. macrocephalus* | 0.0223646 | -0.03930768 | 0.034945546 |
| *S. macrolepis* | 0.002586613 | 0.0056233 | -0.005281302 |
| *S. macrophthalmus* | 0.01921085 | -0.004423179 | 0.011957836 |
| *S. maculatus* | -0.01164108 | -0.003338214 | -0.043022304 |
| *S. maitianheensis* | 0.04058493 | -0.015399253 | -0.004022456 |
| *S. malacopterus* | 0.03369873 | 0.016341474 | 0.002649988 |
| *S. mashanensis* | -0.03299028 | -0.033529198 | -0.005547116 |
| *S. microphthalmus* | -0.02936266 | 0.061223556 | -0.040563771 |
| *S. multipunctatus* | -0.05126049 | 0.014657848 | -0.010821531 |
| *S. oxycephalus* | 0.04140438 | 0.030989746 | -0.036348165 |
| *S. purpureus* | 0.02288238 | -0.045035385 | -0.018448807 |
| *S. qiubeiensis* | 0.03248934 | -0.003880679 | 0.012191068 |
| *S. qujingensis* | 0.000644406 | 0.033555511 | 0.046291334 |
| *S. rhinoceros* | -0.08985807 | 0.03649387 | -0.020510521 |
| *S. ronganensis* | 0.05067357 | -0.004282673 | -0.02013066 |
| *S. tianeensis* | -0.06234116 | 0.02827463 | -0.030564848 |
| *S. tianlinensis* | -0.01905521 | 0.012948686 | -0.00406653 |
| *S. tingi* | 0.01508119 | -0.019031787 | -0.002178747 |
| *S. wumengshanensis* | -0.000171791 | -0.027155405 | 0.003238896 |
| *S. xunlensis* | 0.008420042 | 0.022462118 | 0.047598226 |
| *S. yangzongensis* | 0.02622031 | -0.003887167 | -0.006000758 |
| *S. yimenensis* | -0.01762049 | 0.008265106 | 0.019258774 |
| *S. yishanensis* | 0.007505367 | -0.029610044 | -0.01952727 |
| *S. zhenfengensis* | 0.01938822 | -0.002301361 | 0.003044235 |

